# Monitoring the intestinal magnetic field with optically pumped atomic magnetometers

**DOI:** 10.1101/2022.11.23.517508

**Authors:** Chenxi Sun, Biying Zhao, Teng Wu, Jianwei Zhang, Yuxin Leng, Hong Guo, CREAM Bioelectromagnetism Group

## Abstract

Research has shown the potential of magnetoenterography (MENG) for detecting intestinal diseases noninvasively by superconducting quantum interference devices (SQUIDs). Nevertheless, these devices need to operate under a cytogenetic environment maintained by liquid helium. In this paper, we record the intestinal magnetic field of a rabbit with optically pumped magnetometers (OPMs) at room temperature. It demonstrates that the OPM-based system has sufficient sensitivity to measure the intestinal magnetic fields of the rabbit, and can be potentially developed into a cost-effective and flexible MENG system.

---

Intestinal peristalsis generates electric and magnetic fields through its electrical activities. The fields can reflect real-time functions of the intestinal tracts. Magnetoenterography (MENG) is a technique to measure the magnetic fields produced by electrical currents generated by the smooth muscles in the intestine and the possible induced medium [1]. It has recently garnered interest in clinical medicine for the potential of detecting intestinal diseases noninvasively [2] without serious signal distortion [3, 4], which is superior to electroenterography. Considering the weakness of the magnetic signals, magnetometers with sufficient sensitivity are essential for constructing the measuring system [5]. The conventional systems utilize superconducting quantum interference devices (SQUIDs) and are mostly equipped with magnetic shielding rooms (MSRs), making them expensive to build, maintain and run. In contrast, optically pumped magnetometers (OPMs) can achieve comparable sensitivity with SQUIDs and be cost-effective as well. Nowadays, there is an increasing number of various OPMs being used in noninvasive measurements of cardiac and brain magnetic fields [6, 7, 8, 9].

Our system (shown in Figure 1A) consists of two amplitude-modulated nonlinear magneto-optical rotation (AM-NMOR) magnetometers as a gradiometer and a set of two-layer three-dimensional (3D) Helmholtz coils to actively adjust and stabilize the magnetic field. The system can work at room temperature and in natural magnetic environments, and has been used in measuring bioelectromagnetic signals [6]. With this system, we record the intestinal magnetic fields of the rabbit and assess the feasibility as a replacement and a complementary method to measure MENG signals. The composition and operation of the system are introduced detailly in the supplementary material.

**Figure 1.**
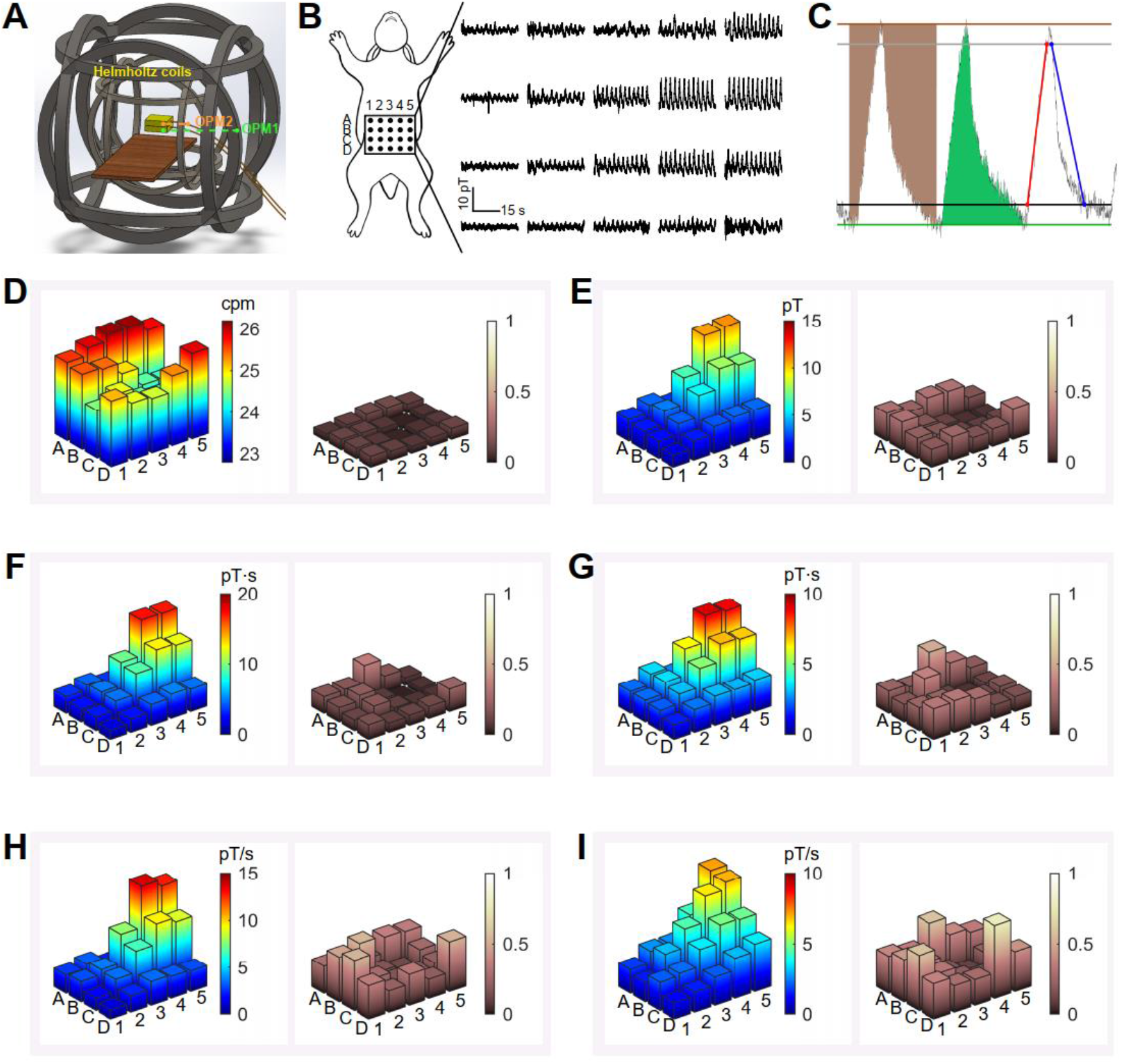
(A) A schematic diagram of our OPM-based MENG system. It consists of a set of two-layer 3D Helmholtz coils to stabilize the environment magnetic field, a gradiometer formed by two AM-NMOR magnetometers to measure the magnetic signals, and a nonmagnetic plate to place the anesthetized laboratory animal. (B) The positions of measuring points and the filtered signals. The A3 point was located at 150 mm down the center of the midpoints of its clavicles, and the distance between the adjoint measuring points was 30 mm. Signals of all the measuring points perform similar frequencies, and those located near the area of the caecum perform much larger amplitudes than others. (C) An illustration of the statistical indicators for quantitively describing the signals. The brown line and the green line show the topline and the baseline, and the brown area and the green area are defined as the AOC and the AUC, respectively. The black line and the gray line show the position of 10% and 90% amplitude, and the slopes of the red line and the blue line correspond to the rising and descending slope, respectively. (D – I) Statistics of the indicators, including frequency (D), amplitude (E), AOC (F), AUC (G), rising slope (H), and descending slope (I), where the left and the right chart of each subfigure show the average value and relative standard deviation of the corresponding indicator, respectively.

The New Zealand white rabbit was anesthetized and then fixed on a nonmagnetic anatomy platform with its venter upwards (details in the supplementary material). The ventral surface was divided into 5 columns, numbered 1 to 5 from right to left of the rabbit, and 4 rows, recorded as A to D from top to bottom (shown in Figure 1B). The A3 point was located at 150 mm down the center of the midpoints of its clavicles, and the distance between the adjoint measuring points was 30 mm. The animal study was reviewed and approved by the Biomedical Ethics Committee of Peking University (ethical approval number: LA2022416).

Figure 1B exhibits the positions of measuring points and the filtered signals. Signals of all the measuring points perform similar frequencies, and those located near positions B4 and B5 perform much larger amplitudes than others, which is consistent with the developed function of the rabbit’s caecum. Figures 1D and 1E present the distribution of frequencies between 23~26 cycles per minute (cpm) and the highest plateau of amplitude above 10 pT. The related measurements by SQUIDs are nearly 3~30 cpm and 1~10 pT [3, 10], showing directly that the OPM-based system has sufficient sensitivity. The increase in the amplitude of our results may be attributed to the shorter distance between the magnetometer and the intestinal tracts in our measuring system, and the variation in the frequency could be caused by the difference in the rabbit fasting time. Individual differences in the testing animals also affect the results.

To describe the signals quantitively, we introduce some statistical indicators (shown in Figure 1C). For each measuring point, we calculate the average value of the signal valleys (peaks) to draw a baseline (topline). For each cycle, the area-under-curve (AUC) is defined as the value of the area between the signal and the baseline, and the area-over-curve (AOC) is that of the area between the signal and the topline. The plateaus of their average values are located in the area near B4 and B5, which is also the basins of their relative standard derivations (shown in Figures 1F and 1G). Besides, for each rising (descending) episode, the two points at 10% and 90% of the amplitude are connected with a line, of which the slope is defined as the rising slope (descending slope). The average values and relative standard derivations of rising and descending slopes in different measuring points exhibit similar distributions as those of the AUC and AOC (shown in Figures 1H and 1I). Furthermore, we also apply the fitness of polynomial functions to every cycle of the measured signals (details in the supplementary material). Notably, current recordings are based on the measurement at discrete time points, in which the large amount of data caused by the high sampling rate places a heavy burden. Instead, the fitted functions contain most of the essential information, not only providing a quantitative method to describe the MENG signals, but also making it possible to decline the cost of signal storage, transmission, procession, and evaluation.

To obtain reams of signals in a limited time and acquire the spatiotemporal correlations of magnetic fields at different positions, an array of sensors is needed. Currently, the multichannel system based on SQUIDs has been developed. It is also possible to establish an array of OPM sensors by utilizing miniaturized vapor cells and compact structures to reduce the space occupied by each channel. As a complement to the fixed measuring system, the system based on the OPM array is of high flexibility and mobility. Getting rid of the Dewar, the position and orientation of each channel can be modified conveniently, leading it to be efficient for obtaining multidirectional signals at different locations as required. Furthermore, it can be potentially developed into wearable systems for different animals and different environments.

## Acknowledgment

We thank Wei Zhang from the National Institute of Metrology of China for assistance with the initial early-stage experimental efforts. We also thank Chen Zhou from the School of Life Sciences of Peking University for his suggestions on rabbit feeding. We also want to thank our funders for making this research possible. A list of funding bodies can be found in the funding section below.

## CREAM Bioelectromagnetism Group

Jingbiao Chen^1^, Shuyuan Chen^1^, Yundi Chang^2^, Xiao Cui^2^, Yudong Ding^1^, Changping Du^1^, Fan Fei^4^, Qinggang Ge^2^, Hong Guo^1^, Junhong Han^1^, Yuxin Leng^2^, Sheng Li^1^, Yike Liang^3^, Meng Liu^1^, Xiyu Liu^1^, Yang Liu^1^, Yuefeng Lu^1^, Yunbin Luo^4^, Yiteng Lv^1^, Xiang Peng^1^, Liang Shen^1^, Mai Shi^2^, Chenxi Sun^1^, Jingyao Suo^4^, Bowen Wang^1^, Haidong Wang^1^, Tianhao Wang^1^, Teng Wu^1^, Tianbo Wu^1^, Wei Xiao^1^, Wei Yan^4^, Dongyi Yang^4^, Xiao Yang^4^, Kaiwen Yi^1^, Jinghua Yu^1^, Chao Zhang^1^, Jianwei Zhang^4^, Pengju Zhang^5^, Qiang Zhang^2^, Shutao Zhang^4^, Biying Zhao^3^, Yixin Zhao^1^, Junhe Zheng^1^, Chao Zhou^1^.

^1^ State Key Laboratory of Advanced Optical Communication Systems and Networks, School of Electronics, and Center for Quantum Information Technology, Peking University, Beijing 100871, China.

^2^ Department of Intensive Care Unit, Peking University Third Hospital, Beijing 100191, China.

^3^ School of Life Sciences, Peking University, Beijing 100871, China.

^4^ School of Physics, Peking University, Beijing 100871, China.

^5^ Faculty of Engineering, University of Bristol, Bristol BS8 1TR, United Kingdom.

## CRediT Authorship Contributions

Chenxi Sun (Conceptualization: Lead; Data curation: Equal; Formal analysis: Equal; Investigation: Lead; Methodology: Equal; Project administration: Supporting; Software: Lead; Validation: Equal; Visualization: Supporting; Writing – original draft: Equal; Writing – review & editing: Equal); Biying Zhao (Conceptualization: Equal; Data curation: Equal; Formal analysis: Equal; Investigation: Equal; Methodology: Supporting; Software: Supporting; Validation: Equal; Visualization: Lead; Writing – original draft: Equal; Writing – review & editing: Equal); Teng Wu (Conceptualization: Supporting; Funding acquisition: Supporting; Project administration: Lead, Supervision: Equal; Writing – review & editing: Equal); Jianwei Zhang (Conceptualization: Equal; Funding acquisition: Supporting; Project administration: Equal, Supervision: Equal; Writing – review & editing: Equal); Yuxin Leng (Conceptualization: Equal; Funding acquisition: Supporting; Project administration: Equal; Resources: Equal; Supervision: Equal; Writing – review & editing: Equal); Hong Guo (Funding acquisition: Lead; Project administration: Equal; Resources: Lead; Supervision: Lead; Writing – review & editing: Equal).

## Conflicts of interest

The authors disclose no conflicts.

## Funding

This work is supported by the National Natural Science Foundation of China (grants nos. 61571018, 61531003, 82172126, and 91436210) and the National Hi-Tech Research and Development (863) Program.

## Notes

### Competing Interest Statement

The authors have declared no competing interest.

## References

[1] S. Somarajan, S. Cassilly, C. Obioha, L.A. Bradshaw, and W.O. Richards, Noninvasive biomagnetic detection of isolated ischemic bowel segments, IEEE Trans. Biomed. Eng. 60, 1677 (2013).

[2] J.P. Wikswo, SQUID magnetometers for biomagnetism and nondestructive testing: important questions and initial answers, IEEE Trans. Appl. Supercond. 5, 74 (1995).

[3] L.A. Bradshaw, S.H. Allos, J.P. Wikswo, and W.O. Richards, Correlation and comparison of magnetic and electric detection of small intestinal electrical activity, Am. J. Physiol. Gastrointest. 272, G1159 (1997).

[4] L.A. Bradshaw, W.O. Richards, and J.P. Wikswo, Volume conductor effects on the spatial resolution of magnetic fields and electric potentials from gastrointestinal electrical activity, Med. Biol. Eng. Comput. 39, 35 (2001).

[5] J. Golzarian, D. Staton, J. Wikswo, R.N. Friedman, W. Richards, First biomagnetic measurements of intestinal basic electrical rhythms (BER) *in vivo* using a high-resolution magnetometer, Gastroenterology 103, 1385 (1992).

[6] C. Sun, W. Xiao, Y. Ding, R. Zhang, M. Liu, T. Wu, X. Peng, J. Chen, and H. Guo, Recording the distribution of cardiac magnetic fields in unshielded earth’s field, in 2021 Joint Conference of the European Frequency and Time Forum and IEEE International Frequency Control Symposium (EFTF/IFCS) (IEEE, 2021) pp. 1–3, and references therein.

[7] A.U. Kowalczyk, Y. Bezsudnova, O. Jensen, and G. Barontini, Detection of human auditory evoked brain signals with a resilient nonlinear optically pumped magnetometer, NeuroImage 226, 117497 (2021).

[8] G. Bison, N. Castagna, A. Hofer, P. Knowles, J.-L. Schenker, M. Kasprzak, H. Saudan, and A. Weis, A room temperature 19-channel magnetic field mapping device for cardiac signals, Appl. Phys. Lett. 95, 173701 (2009).

[9] E. Boto, N. Holmes, J. Leggett, G. Roberts, V. Shah, S.S. Meyer, L.D. Muñoz, K.J. Mullinger, T.M. Tierney, S. Bestmann, et al., Moving magnetoencephalography towards real-world applications with a wearable system, Nature 555, 657 (2018).

[10] L.A. Bradshaw, C.L. Garrard, J.P. Wikswo, and W.O. Richards., Transabdominal magnetic recording of small bowel ischemia in anesthetized rabbits, Gastroenterology 107, 4 (1994).

