## Supplementary Materials for "Monitoring the intestinal magnetic field with optically pumped atomic magnetometers"

**Supplementary Material**

*The Measurement System*

Our amplitude-modulated nonlinear magneto-optical rotation (AM-NMOR) magnetometers are self-made and have a size of $5\ \text{cm} \times 24\ \text{cm} \times 27\ \text{cm}$, mounted in a three-dimensional printed structure. All the components are non-magnetic, including a cylinder vapor cell and optical elements consisting of mirrors, polarizers, wave plates, prisms, etc. The diameter and the length of the vapor cell are both 25 mm. The basic structure, first proposed in the 1960s [1], is a pump-probe scheme with an atomic vapor cell. The self-made cesium vapor cell in our magnetometer is coated with paraffin to increase the relaxation time of the coherent state of the cesium atoms [2] and had a typical magnetic-resonance linewidth of ~7 Hz at room temperature. A circularly-polarized pump laser transmits the vapor cell to polarize the cesium atoms and form a macroscopic magnetic moment precessing at the Larmor frequency [3]. The pump laser is generated and locked to the frequency of the cesium D1 transition line by the laser generator (Toptica, DL pro) with its controller, and modulated on and off at the Larmor frequency with a duty cycle of 20% by a waveform generator (Keysight, 33522B) and an acoustic-optical modulator (AOM, AA Opto Electronic, MT350-B120A0,12-800). A continuous linearly-polarized probe laser transmits the vapor cell perpendicular to the pump laser to detect the evolution of the atomic magnetic moment [3]. The probe laser is generated and locked to the frequency positively detuned by ~400 MHz from the D2 transition line. The frequencies of both the pump and probe lasers are actively stabilized with dichroic atomic vapor laser lock modules. The probe beam is detected by a balanced photodetector (Thorlabs, PDB210A/M) and demodulated with a lock-in amplifier (LIA, Stanford Research Systems, SR865A).

To prevent the distortion of signals, we apply the closed-loop method to expand the bandwidth of the gradiometers [4]. The modulation frequency of the pump laser is adjusted by an analog proportional integral derivative (PID) controller (Stanford Research Systems, SIM960) to track the Larmor frequency under the ambient magnetic field. The signals are sensed as the change of environment magnetic fields by the probe light, demodulated by the LIA, and feedback to the modulation frequency of the pump laser by the PID controller. The value of the magnetic field measured by the magnetometer is equal to the modulation frequency of the pump laser divided by the gyromagnetic ratio of the cesium atom.

The outer layer of the Helmholtz coils shown in Figure 1A is a set of three orthogonal pairs of coils with 3-meter diameters for adjusting the direction and strength of the bias magnetic field. The horizontal components of the bias magnetic field are adjusted to zero with a fluxgate magnetometer, and the vertical component is set to 20,000 nT to avoid the decreasing of the sensitivity of the gradiometer due to the nonlinear Zeeman effect.

The active magnetic-field stabilization method is applied to suppress the environment magnetic-field noise. The magnetic-field stabilization system consists of a magnetic-field generator, a magnetic-field sensor, and a feedback controller [5]. In our system shown in Figure 1A, the top OPM works as the magnetic-field sensor to monitor the environment noise, and the pair of coils along the vertical direction of the inner layer of 3D Helmholtz coils works as the magnetic field generator. Considering our OPM sensors are not sensitive to magnetic-field noise along the horizontal directions, only the magnetic field in the vertical direction was actively stabilized. An analog PID controller (Stanford Research Systems, SIM960) works as the feedback controller to adjust the current in the vertical Helmholtz coil via a voltage-controlled current (VOC) source (Stanford Research Systems, LDC501), to compensate for the fluctuations of the magnetic field around the vapor cell of the top OPM to a fixed setpoint.

*Data Acquisition and Signal Analysis*

Due to the limited use time of the laboratory, we have only measured the distribution of MENG of one rabbit subject. The rabbit subject is a female adult New Zealand white rabbit, with a weight of 4.3 kg. It is anesthetized with sodium thiopental (20 mg/kg) and then fixed on the nonmagnetic anatomy platform made up of wood and plastic. The prototype gradiometer is moved in S-shape to measure the signal at the positions shown in Figure 1B. Each position is measured for 30 seconds, from A1 to D5 by turns. The digital MENG signals are collected at a sampling rate of 40 kSa/s with a data acquisition card (National Instruments, USB6363) and a LabVIEW-based data acquisition system. All data processing and signal analysis are realized in MATLAB (MathWorks, Natick, MA). The recorded raw MENG signals are filtered to remove the power frequency noise and harmonics with Butterworth band-stop filters and then downsampled to 40 Hz for analysis. The low-frequency fluctuations of the signals are also removed by subtracting the 5-second moving average.

*Fitness of Polynomial Functions*

Based on the results of our measurement, we apply the fitness of polynomial functions to the collected data in the rising and descending episode of every cycle. Considering the features of the shapes and turning points in every cycle, we choose the cubic polynomial function and use the ordinary least squares regression to estimate the coefficients. The four coefficients in descending order of power for the obtained function of the rising (descending) episode are recorded as parameters $a$ ($k$), $b$ ($l$), $c$ ($m$), and $d$ ($n$), respectively. Figure S1 shows the statistics of these parameters in six measuring positions.

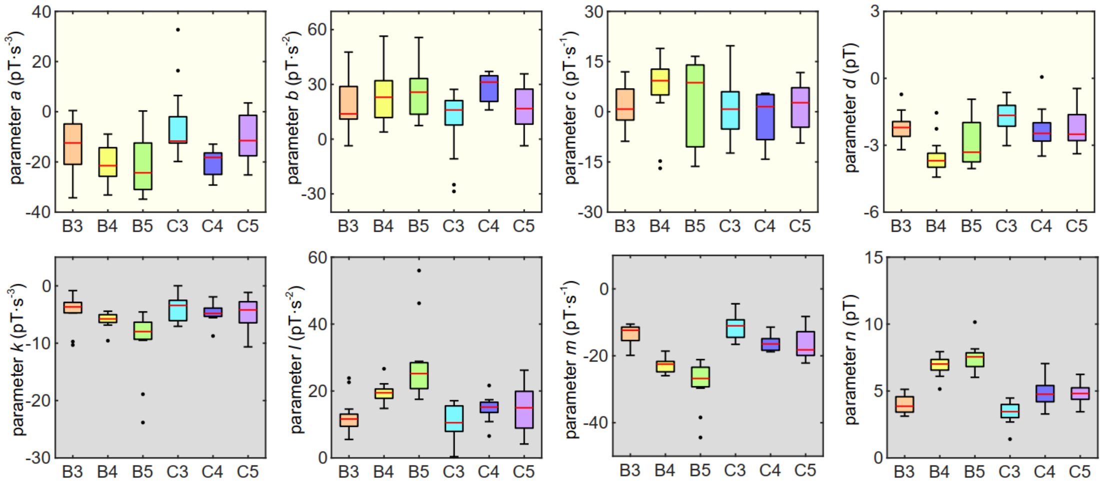


**Figure S1** Statistics of the parameters in six measuring points. Boxplots for the coefficients in descending order of power for the obtained function of the rising episode and the descending episode are shown in the first and second row, respectively. The red lines represent the median values. The lower and upper edges of the boxes represent the lower quartiles and the upper quartiles, respectively. The lower and upper edges of the vertical lines represent the maximum and the minimum values of the data. The single points show the outliers.

*References of Supplementary Material*


[1] W.E. Bell and A.L. Bloom, Optically driven spin precession, Phys. Rev. Lett. **6**, 280 (1961).

[2] J. Dupont-Roc, S. Haroche, and C. Cohen-Tannoudji, Detection of very weak magnetic fields ($10^{-9}$ gauss) by $^{87}$Rb zero-field level crossing resonances, Phys. Lett. A **28**, 638 (1969).

[3] D.F.J. Kimball, S. Pustelny, V.V. Yashchuk, and D. Budker, in *Optical Magnetometry*, edited by D. Budker and D.F.J. Kimball (Cambridge University Press, Cambridge, 2013), pp. 104-124.

[4] R. Zhang, W. Xiao, Y. Ding, Y. Feng, X. Peng, L. Shen, C. Sun, T. Wu, Y. Wu, Y. Yang, Z. Zheng, X. Zhang, J. Chen, and H. Guo, Recording brain activities in unshielded earth's field with optically pumped atomic magnetometers, Sci. Adv. **6**, eaba8792 (2020).

[5] R. Zhang, Y. Ding, Y. Yang, Z. Zheng, J. Chen, X. Peng, T. Wu, and H. Guo, Active magnetic-field stabilization with atomic magnetometer, Sensors **20**, 4241 (2020).
